# Nucleus accumbens astrocytes bidirectionally modulate social behavior

**DOI:** 10.1101/2024.09.18.613653

**Authors:** Jonathan W VanRyzin, Kathryn J Reissner

**Affiliations:** Department of Psychology and Neuroscience, University of North Carolina at Chapel Hill, Chapel Hill, NC 27514

**Keywords:** astrocytes, nucleus accumbens, DREADDs, hPMCA, social behavior

## Abstract

Social behaviors are critical for survival and fitness of a species, and maladaptive social behaviors are frequently associated with neurodevelopmental and psychiatric disorders. As such, the neural circuits and cellular mechanisms driving social behaviors inform critical processes contributing to both health and disease. In particular, the nucleus accumbens (NAc) is a key hub for the integration of both social and non-social information required for successful social interactions and reward motivated behaviors. While astrocytes within the NAc have a recognized role in modulating neural activity, their influence over social behavior is yet undefined. To address this question, we manipulated NAc astrocyte signaling and determined effects on social interactions. NAc core astrocytes bidirectionally influenced social behavior in rats; agonism of astrocyte-specific hM3D(Gq) DREADD receptors increased social interaction time in the social interaction test and increased social preference in the 3-chamber test. Conversely, decreasing intracellular calcium signaling in astrocytes with viral expression of hPMCA reduced both social interaction and social preference in these tests. These results suggest that NAc astrocytes actively participate in the regulation of social behavior and highlight a putative role for astrocytes in disorders characterized by social dysfunction.

## INTRODUCTION

The appropriate expression of social behaviors is essential for the survival and reproduction of a species. To regulate these behaviors, the brain must process a variety of signals, including social, affective, motivational, and reward-related information (O’Connell & Hofmann, 2011; Stanley & Adolphs, 2013). The nucleus accumbens (NAc) plays a key role in integrating this information to shape social behavior, processing neural activity from numerous brain regions and varied neurotransmitters and neuromodulators (Kohls et al., 2013; Dolen et al., 2013; Manduca et al., 2016; Schmidt et al., 2019; Shan et al., 2022; Le Merrer et al., 2024).

Within the NAc, astrocytes are uniquely poised to both detect, and respond to, neural inputs via a diversity of neurotransmitter receptors (Perea et al., 2009). Stimulation of astrocytic neurotransmitter receptors by synaptic activity induces intracellular calcium signaling cascades, leading to the release of gliotransmitters that can reciprocally modulate synaptic activity and plasticity (Araque 2009; Dallerac et al., 2013; Harada et al., 2016). Importantly, recent evidence implicates astrocyte control of neurotransmission as important for social behaviors: cortical astrocytes regulate social hierarchy in male mice by influencing excitatory/inhibitory balance (Noh et al., 2023) and social isolation dysregulates astrocytic GABA release in the hippocampus (Cheng et al., 2023). However, little is known regarding whether or how NAc astrocytes contribute to social interactions. Thus, we sought to determine to extent to which NAc astrocytes influenced the expression of social behaviors.

Here, we show that NAc core astrocytes bidirectionally modulate the expression of social behaviors. Chemogenetic Gq stimulation of astrocytes increased social interaction time and social preference, while conversely, decreasing astrocyte intracellular calcium signaling decreased both social interaction time and social preference. Together, these results demonstrate a role for NAc core astrocyte Ca2+ dynamics in the basal expression of social behavior.

## MATERIALS AND METHODS

### Animals

Adult male Sprague Dawley rats (225-250 g) were purchased from Envigo and were kept on a 12:12 h reverse light/dark cycle with ad libitum food and water. Experimental animals were allowed to acclimate for at least 1 week upon arrival and were single housed following surgical procedures, while stimulus animals were housed in groups of 2-3 rats for the duration of these studies. All animal care and procedures were performed in accordance with the National Institutes of Health *Guide for the Care and Use of Laboratory Animals* and were approved by the Institutional Animal Care and Use Committee at the University of North Carolina at Chapel Hill.

### Surgical procedures

Animals were anesthetized with ketamine (100 mg/kg) and xylazine (7 mg/kg) and head-fixed in a stereotaxic apparatus (Kopf Instruments). Viral vectors were delivered bilaterally using a 24 gauge syringe (Hamilton) at a rate of 100 nl/min and allowed to diffuse for an additional 10 min following infusion. The nucleus accumbens (NAc) core was targeted using the following surgical coordinates:

NAc core (+1.5 AP, ±2.6 ML, -7.2 DV, 6° angle). The following viral constructs were used in these studies:

AAV5-GfaABC1D-Lck-mCherry (1 µl/hemisphere; 6 × 10^12^ viral particles/ml; packaged at University of North Carolina (UNC) Viral Vector Core). Plasmid was a kind gift of Joshua Jackson at Drexel University.

AAV5-GfaABC1D-hM3D(Gq)-mCherry (1 µl/hemisphere; 1.1 × 10^13^ viral particles/ml; Addgene plasmid #92284; packaged at UNC Viral Vector Core) (Chai et al., 2017).

AAV5-GfaABC1D-mCherry-hPMCA2w/b (1 µl/hemisphere; 1.0 × 10^13^ viral particles/ml; Addgene #111568-AAV5) (Yu et al., 2018).

Animals were allowed to recover for at least 3 weeks before behavioral analysis to ensure proper viral expression.

### Behavioral assays

All behavioral testing was performed during the dark phase of the light-dark cycle under red light illumination. All behavioral tests were performed, video recorded, and scored offline by an experimenter blind to experimental manipulation. For DREADD-activation experiments, animals were randomly assigned to treatment groups and counterbalanced across test days. Prior to behavioral testing, DREADD-expressing animals were allowed to acclimate 30 min, after which, they received either clozapine N-oxide (CNO) (5 mg/kg; i.p.; HelloBio #HB6149) or saline (i.p.) injections. Animals then remained in the testing room for an additional 30 min prior to the start of behavioral assessment. All behavioral testing was performed on consecutive days.

#### Social Interaction Test

Experimental animals were placed in a novel enclosure (24 × 18 × 12 in) with TEK-Fresh cellulose bedding (Harland Laboratories). At the start of the experiment, a novel age- and sex-matched, previously group-housed stimulus animal was introduced to the enclosure and allowed to freely interact for 5 min. The experimental animal’s behavior was assessed to quantify the time spent engaging in active social investigation (sniffing, licking, grooming, etc) of the stimulus animal.

#### 3 Chamber Social Preference Test

The day before testing, animals were habituated to a 3-chamber arena (54 × 36 × 12 in) for 10 min. At the start of testing, a novel age- and sex-matched stimulus animal was placed under an enclosure in either the right or left side of the arena, while the opposite side enclosure was left empty. The position of the stimulus animal was counterbalanced across experimental animals to avoid any confounds due to inherent side preference. The experimental animal was then placed into the center of the arena and allowed to freely explore for 5 min. The experimental animal’s behavior was assessed to quantify the time spent actively investigating the social and empty enclosures, as well as entries into the social and empty sides of the arena. A Social Preference Index was calculated as (Social Investigation Time)/(Social Investigation Time + Empty Investigation Time).

### Immunohistochemistry

Animals were anesthetized with sodium pentobarbital and transcardially perfused with phosphate buffered saline (PBS; 0.1M, pH 7.4) followed by 4% paraformaldehyde (PFA, 4% in PBS, pH 7.2).

Brains were extracted, postfixed in 4% PFA for 24 hours at 4°C, then kept in 30% sucrose solution at 4°C until fully submerged. Coronal sections were cut at a thickness of 45 µm using a cryostat (Leica Biosystems) and stored in cyroprotectant solution until processing.

Tissue sections were mounted onto slides and washed with PBS and blocked with 5% bovine serum albumin (BSA) in PBS + 0.4% Triton X-100 (PBS-T) for 1 h. Slides were incubated in primary antibody solution (2.5% BSA in PBS-T) overnight at RT. The next day, slides were incubated in secondary antibody solution (2.5% BSA in PBS-T) for 2 h, stained with Hoescht 33342 (1:2000; Invitrogen #H3570), and coverslipped with ProLong Diamond Antifade Mountant (Thermo Fisher Scientific). The following primary antibodies were used in these studies: Mouse anti-NeuN (1:1000; EMD Millipore #MAB377), Rabbit anti-mCherry (1:1000; Rockland #600-401-379). The following secondary antibodies were used in these studies: Alexa Fluor (all 1:500; Thermo Fisher Scientific): Donkey anti-rabbit 594, Donkey anti-mouse 647.

### Image acquisition

Confocal fluorescence images were acquired using a Zeiss LSM 800 equipped with 405, 488, 561, and 640 lasers and a 20x (0.8 NA) and 63x (1.4 NA) oil-immersion objective. Images were acquired using between 2.5-5.0 μm z-step for 20x images and 0.5 μm z-step for 63x images and processed to correct for background signal in ImageJ.

### Statistical analysis

Statistical analysis was performed using R (R Core Team, 2021; version 4.1.0). All values are shown as the mean ± SEM. Comparisons between two experimental groups were performed using a two-tailed Welch’s t-test or paired t-test for between or within subject designs, respectively. A p value of < 0.05 was used to determine significance. Additional statistical details of each experiment (such as n, t value, etc) can be found in the figure legends and in the text.

## RESULTS

### Astrocytes bidirectionally modulate social behavior

To determine how astrocytes within the nucleus accumbens (NAc) core may regulate social behavior, we first used astrocyte-specific hM3D(Gq) DREADDs to chemogenetically alter astrocyte function, which have been previously shown to induce intracellular calcium signaling and gliotransmitter release from astrocytes (Chai et al., 2017; Durkee et al., 2019). We bilaterally infused AAV5-GfaABC1D-hM3D(Gq)-mCherry into the NAc core of adult male rats and allowed to express for 3 weeks before behavioral testing (Figure 1A). hM3d(Gq)-mCherry expression was mostly restricted to the NAc core and showed highly branched and diffuse expression typical of astrocytes. Importantly, hM3D(Gq)-mCherry expression did not colocalize with NeuN, a neuronal cell marker (Figure 1B). To assess social behavior, animals underwent behavioral assessment in two common assays of social behavior: the social interaction test and the social preference test (Figure 1C). For each test, a novel age- and sex-matched conspecific was used as a stimulus animal. Animals were tested in each paradigm twice, receiving either saline or CNO (to stimulate hM3D(Gq)) in a counterbalanced design, and behavior was analyzed as a within-subjects design across test days.

**Figure 1.**
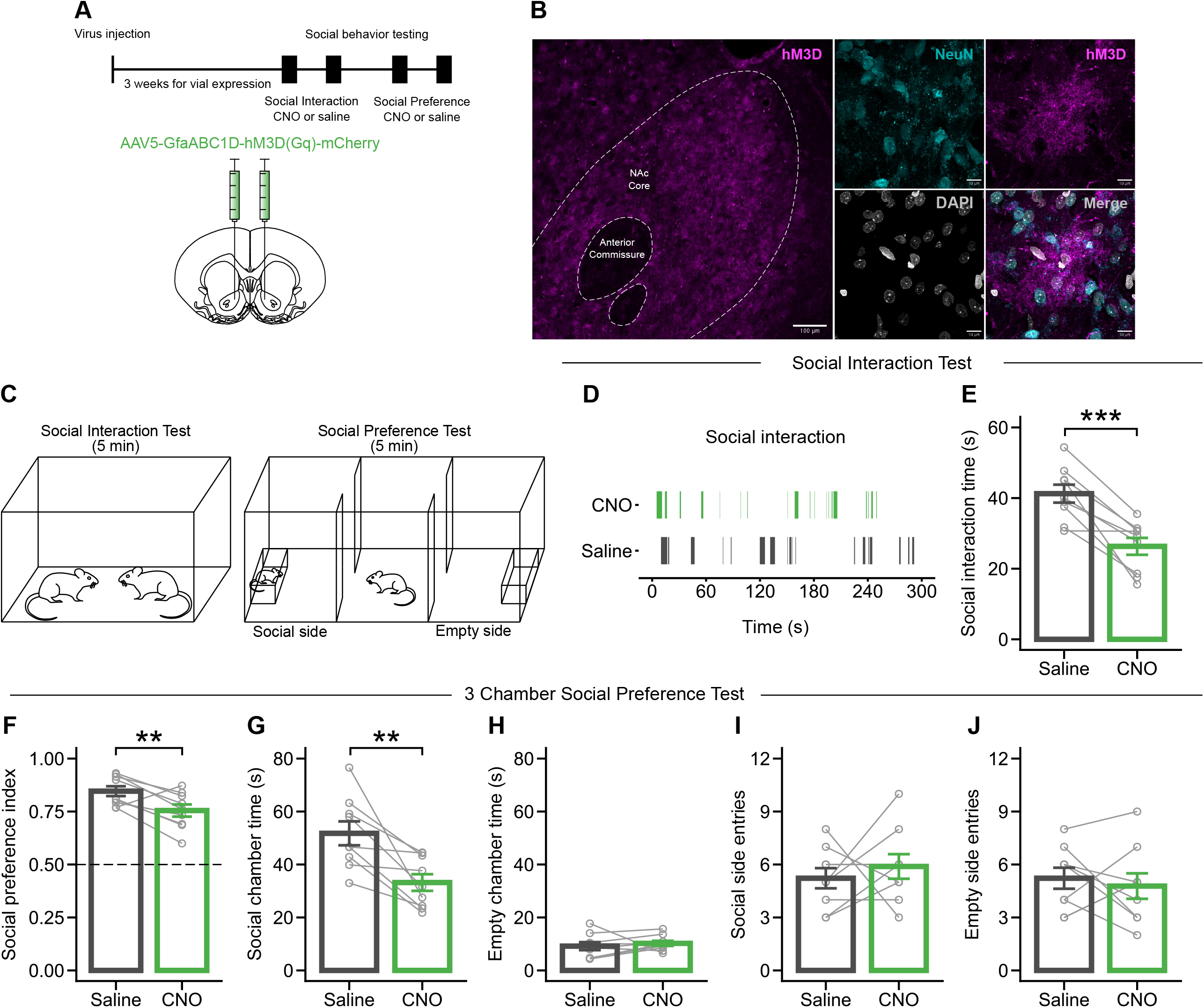
Chemogenetic stimulation of astrocyte-specific hM3D(Gq) in the nucleus accumbens core decreases social interaction and social preference. (A) Experimental timeline. AAV5-GfaABC1D-hM3D(Gq)-mCherry was injected into the NAc core 3 weeks prior to behavioral testing. n = 9 rats. (B) Viral expression of GfaABC1D-hM3D was largely restricted to the NAc core (left). High magnification imaging shows hM3D expression highlighting classic astrocyte morphology and did not colocalize with the neuron marker, NeuN, suggesting astrocyte-specific expression (right). Scale bar represents 100 µm (left) and 10 µm (right). (C) Depiction of the social interaction and social preference tests. (D) Representative ethogram from a single animal in the social interaction test during the CNO and saline trials. (E) Chemogenetic stimulation of hM3D with CNO decreased social interaction time in the social interaction test and decreased preference index (F) and time spent interacting with the social chamber (G), but did not affect time spent interacting with the empty chamber (H) or the number of entries into either the social (I) or empty (J) side of the social preference test. Bars represent the mean ± SEM. Open circles represent data points for each animal, grey bars indicate values for an individual animal across trials. NAc, nucleus accumbens. **p < 0.01, ***p < 0.001.

In the social interaction test, CNO treatment decreased social interaction time compared to saline treatment (Figure 1D, 1E; t(8) = 5.5733, p < 0.001). CNO treatment also decreased the preference index in the social preference test (Figure 1F; t(8) = 3.3852, p = 0.009). This effect was largely driven by a corresponding decrease in the time spent investigating the social chamber that housed the novel stimulus animal (Figure 1G; t(8) = 3.861, p = 0.004), with no change in time spent investigating the empty chamber between CNO- and saline-treatment days (Figure 1H). Moreover, we found no change in the number of times animals entered either the social or empty side of the arena (Figure 1I, 1J), suggesting that the observed decreases in social behavior were not due to changes in locomotor or exploratory behavior.

To assess whether CNO treatment had any effect on social behavior independent of DREADD activity, we repeated the same experimental paradigm in animals that were bilaterally injected with a control virus, AAV5-GfaABC1D-mCherry, into the NAc core (Figure S1A, S1C). Following 3 weeks of viral incubation, mCherry expression was similarly localized to astrocytes in the NAc core (Figure S1B). As expected, we found no difference in social interaction time between saline or CNO trials in the social interaction test (Figure S1D, S1E), and found no effect of CNO treatment on preference index (Figure S1F), social or empty chamber time (Figure S1G, S1H), and social or empty side entries (Figure S1I, S1J). Together, these data show that the observed decreases in social behavior are due to DREADD stimulation-induced changes in NAc astrocyte function.

Given that hM3D stimulation of astrocytes has been shown to increase intracellular calcium signaling and alter astrocyte function, we next sought to diminish calcium signaling within astrocytes and determine the impact on social behavior. To this end, we bilaterally infused AAV5-GfaABC1D-mCherry-hPMCA2w/b, which encodes for a constitutively active calcium extrusion pump and decreases the frequency and amplitude of calcium signaling within astrocytes (Yu et al., 2018; Yu et al., 2021), or AAV5-GfaABC1D-Lck-mCherry, into the NAc core (Figure 2A). hPMCA expression was widespread within the NAc core and highly specific to astrocytes (Figure 2B). Animals then underwent a single session of both social interaction and social preference tests. Interestingly, hPMCA expression significantly increased social interaction time in the social interaction test, compared to controls (Figure 2D, 2E; t(16.61) = -2.7401, p = 0.014). Similarly, there was increased preference index (Figure 2F; t(17.304) = -5.0315, p < 0.001), which was driven by a trending but non-significant increase in social chamber time (Figure 2G; t(12.26) = -1.8557, p = 0.087) and a decrease in empty chamber time (Figure 2H; t(13.58) = 4.3128, p < 0.001) in hPMCA-expressing animals compared to mCherry-expressing controls. We found no difference in social or empty side entries between groups (Figure 2I, 2J).

**Figure 2.**
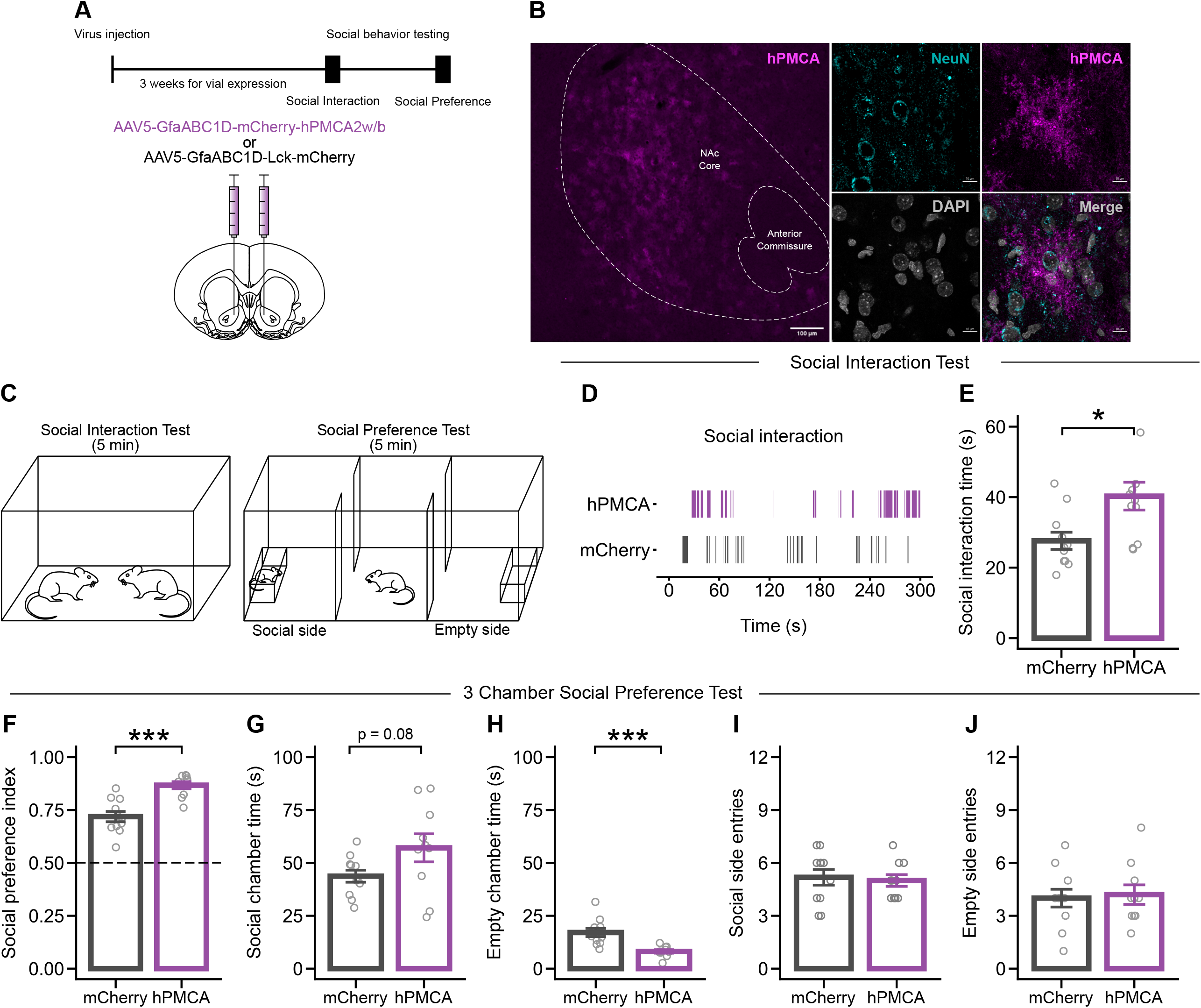
Decreasing astrocyte calcium signaling in the nucleus accumbens core increases social interaction and social preference. (A) Experimental timeline. AAV5-GfaABC1D-mCherry-hPMCA2w/b was injected into the NAc core 3 weeks prior to behavioral testing. n = 11 rats. (B) Viral expression of GfaABC1D-hPMCA2w/b was largely restricted to the NAc core (left). High magnification imaging shows hPMCA expression highlighting classic astrocyte morphology and did not colocalize with the neuron marker, NeuN, suggesting astrocyte-specific expression (right). Scale bar represents 100 µm (left) and 10 µm (right). (C) Depiction of the social interaction and social preference tests. (D) Representative ethogram from hPMCA or mCherry-expressing animals during the social interaction test. (E) hPMCA expression increased social interaction time in the social interaction test. (F) hPMCA expression increased preference index, did not affect time spent interacting with the social chamber (G), and decreased time spent interacting with the empty chamber (H). hPMCA expression did not affect the number of entries into either the social (I) or empty (J) side of the social preference test. Bars represent the mean ± SEM. Open circles represent data points for each animal. NAc, nucleus accumbens. *p <0.05, ***p < 0.001.

## DISCUSSION

The current study shows that NAc core astrocytes are essential for the proper expression of social behavior. By manipulating intracellular astrocytic calcium, we show that astrocytes bidirectionally modulate social interactions and social preference. These findings highlight astrocytes within the NAc core as a possible convergence point for “tuning” social behavior.

The neurobiology of social behavior is complex, requiring the integration of social, affective, motivational, and reward information. To this end, it has been proposed that the processing of social information is distributed across a network of brain regions referred to as the social behavior network (Newman, 1999; Goodson, 2005). In this framework, the NAc acts as a hub which interfaces the mesolimbic reward system with the processing of social stimuli (O’Connell & Hofmann, 2011; Xu et al., 2021; Walsh et al., 2023). Neural activity within the NAc has been implicated in human (Kohls et al., 2013; Schmidt et al., 2019) and rodent studies, the latter demonstrating that NAc activity is important for social approach, social dominance, and social reward (Dolen et al., 2013; Gunaydin et al., 2014; Ferri et al., 2016; Manduca et al., 2016; Shan et al., 2022; Le Merrer et al., 2024).

Astrocytes can detect and regulate neural transmission (Perea et al., 2009; Nedergaard & Verkhratsky, 2012). Detection of synaptic activity via membrane-bound neurotransmitter receptors initiates intracellular calcium signaling cascades that lead to the release of gliotransmitters that reciprocally modulate synaptic activity and plasticity (Araque, 2009; Dallerac et al., 2013; Harada et al., 2016). Commonly, gliotransmitter release induces synaptic depression of presynaptic inputs, largely through agonism of inhibitory presynaptic receptors (Corkrum et al., 2020; Kofuji & Araque, 2021; Ortinski et al., 2022). Our data show that altering intracellular calcium signaling within NAc core astrocytes can mediate social behavior: decreasing intracellular calcium via hPMCA expression (Yu et al., 2018) increases social interaction and preference, while increasing calcium via hM3D(Gq) DREADD activation (Chai et al., 2017) decreases social interaction and preference. We interpret the ability of astrocytes to bidirectionally modulate social behavior as evidence that astrocytes influence neural signaling within the NAc core to affect behavioral output. Activating astrocyte Gq signaling, and presumably inducing gliotransmitter release, decreases social behavior, while reducing the ability of astrocytes to depress neurotransmission is permissive for increased social behavior. It remains to be determined whether astrocytes regulate specific neural circuits within the NAc that mediate social behavior, or whether astrocytes broadly govern neural activity to impact behavior.

Astrocytes have been previously implicated in regulating a range of behaviors, including learning and memory, locomotor activity, anxiety-like, and social behaviors among others (Erta et al., 2015; Adamsky et al., 2018; Gomez et al., 2019; Wang et al., 2021; Cho et al., 2022; Noh et al., 2023; Meadows et al., 2024; reviewed in Kofuji & Araque, 2021; Barnett et al., 2023). Our findings support these conclusions by showing that astrocytes can effectively act to “maintain” social behavior within an optimal range. Future studies should investigate how psychiatric disorders or behavioral paradigms that produce alterations in social behavior also alter astrocytes, as they may be a common denominator by which alterations in behavior occur.

## FIGURE LEGENDS

**Supplementary Figure 1.**
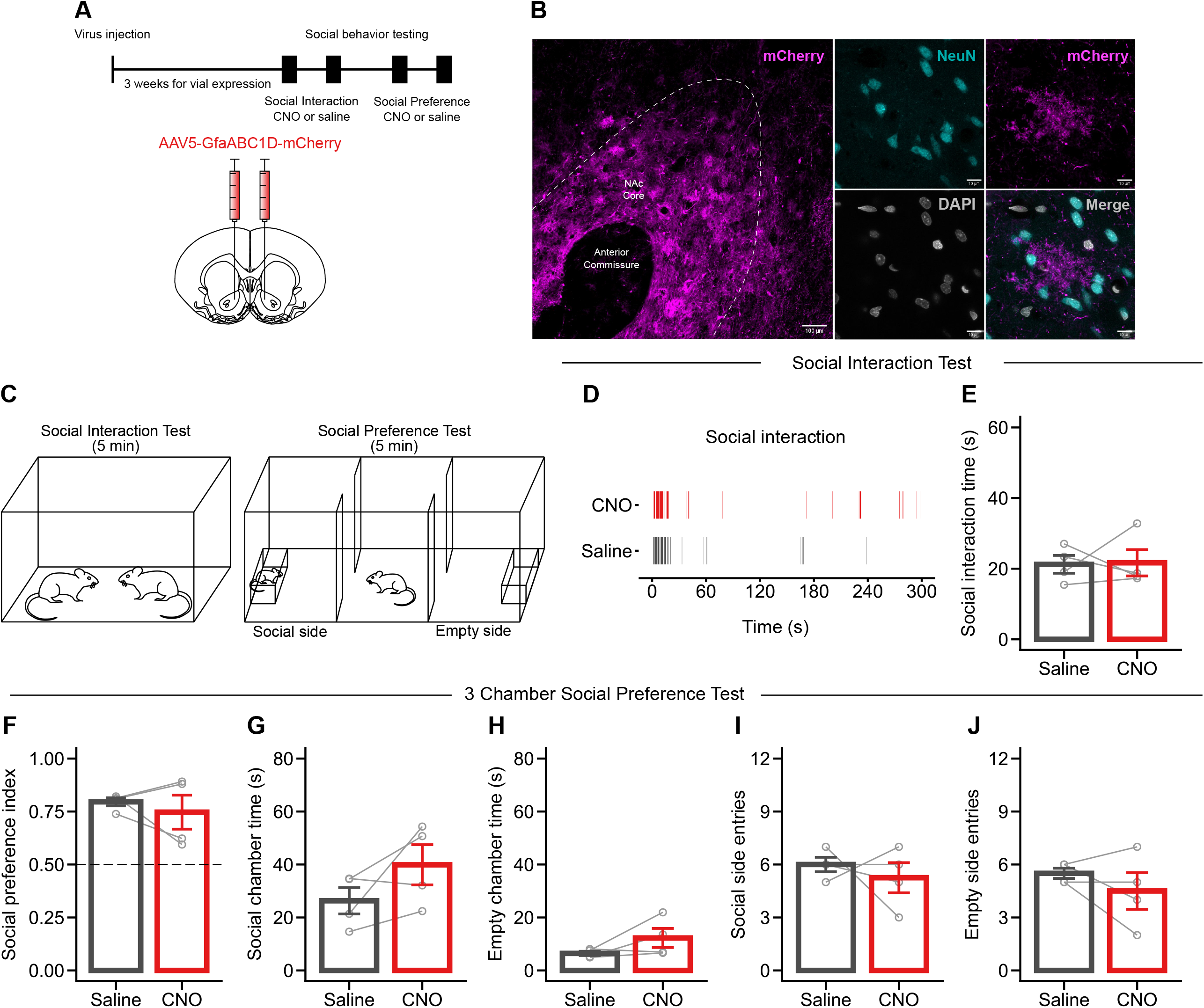
CNO treatment alone did not affect social behavior. (A) Experimental timeline. AAV5-GfaABC1D-mCherry was injected into the NAc core 3 weeks prior to behavioral testing. n = 4 rats. (B) Viral expression of GfaABC1D-mCherry was largely restricted to the NAc core (left). High magnification imaging shows mCherry expression highlighting classic astrocyte morphology and did not colocalize with the neuron marker, NeuN, suggesting astrocyte-specific expression (right). Scale bar represents 100 µm (left) and 10 µm (right). (C) Depiction of the social interaction and social preference tests. (D) Representative ethogram from a single animal in the social interaction test during the CNO and saline trials. (E) CNO treatment did not affect social interaction time in the social interaction test or preference index (F), time spent interacting with the social chamber (G), time spent interacting with the empty chamber (H), entries into either the social (I) or empty (J) side of the social preference test. Bars represent the mean ± SEM. Open circles represent data points for each animal, grey bars indicate values for an individual animal across trials. NAc, nucleus accumbens. **p < 0.01, ***p < 0.001.

## REFERENCES

1. Adamsky A, Kol A, Kreisel T, Doron A, Ozeri-Engelhard N, Melcer T, Refaeli R, Horn H, Regev L, Groysman M, London M, Goshen I. Astrocytic activation generates de novo neuronal potentiation and memory enhancement. Cell. 2018; 174:59–71.

2. Araque A. Astrocytes process synaptic information. Neuron Glia Biol. 2009; 4:3–10.

3. Barnett D, Bohmbach K, Grelot V, Charlet A, Dallérac G, Ju YH, Nagai J, Orr AG. Astrocytes as drivers and disruptors of behavior: new advances in basic mechanisms and therapeutic targeting. J Neurosci. 2023; 43:7463–7471.

4. Chai H, Diaz-Castro B, Shigetomi E, Monte E, Octeau JC, Yu X, Cohn W, Rajendran PS, Vondriska TM, Whitelegge JP, Coppola G, Khakh BS. Neural circuit-specialized astrocytes: transcriptomic, proteomic, morphological and functional evidence. Neuron. 2017; 95:531–549.

5. Cheng Y-T, Woo J, Luna-Figueroa E, Maleki E, Harmanci AS, Deenen B. Social deprivation induces astrocytic TRPA1-GABA suppression of hippocampal circuits. Neuron. 111:1301-1315.e5.

6. Cho H-W, Noh K, Lee BH, Barcelon E, Jun SB, Park HY, Lee SJ. Hippocampal astrocytes modulate anxiety-like behavior. Nat Commun. 2022; 13:6536.

7. Corkrum M, Covelo A, Lines J, Bellocchio L, Pisansky M, Loke K, Quintana R, Rothwell PE, Lujan R, Marsicano G, Martin ED, Thomas MJ, Kofuji P, Araque A. Dopamine-evoked synaptic regulation in the nucleus accumbens requires astrocyte activity. Neuron. 2020; 105:1036–1047.

8. Dallerac G, Chever O, Rouach N. How do astrocytes shape synaptic transmission? Insights from electrophysiology. Front Cell Neurosci. 2013; 7:159.

9. Dolen G, Darvishzadeh A, Huang KW, Malenka RC. Social reward requires coordinated activity of nucleus accumbens oxytocin and serotonin. Nature. 2013; 501:179–184.

10. Durkee CA, Covelo A, Lines J, Kofuji P, Aguilar J, Araque A. Gi/o protein coupled receptors inhibit neurons but activate astrocytes and stimulate gliotransmission. Glia. 2019; 67(6):1076–93.

11. Erta M, Giralt M, Esposito FL, Fernandez-Gayol O, Hidalgo J. Astrocytic IL-6 mediates locomotor activity, exploration, anxiety, learning and social behavior. Horm Behav. 2015; 73:64–74.

12. Ferri SL, Kreibich AS, Torre M, Piccoli CT, Dow H, Pallathra AA, Li H, Bilker WB, Gur RC, Abel T, Brodkin ES. Activation of basolateral amygdala in juvenile C57BL/6J mice during social approach behavior. Neuroscience. 2016; 335:184–194.

13. Goodson JL. The vertebrate social behavior network: evolutionary themes and variations. Horm Behav. 2005; 38:11–22.

14. Gomez JA, Perkins JM, Beaudoin GM, Cook NB, Quraishi SA, Szoeke EA, Thangamani K, Tschumi CW, Wanat MJ, Maroof AM, Beckstead MJ, Rosenberg PA, Paladini CA. Ventral tegmental area astrocytes orchestrate avoidance and approach behavior. Nat Commun. 2019; 10:1455.

15. Gunaydin LA, Grosenick L, Finkelstein JC, Kauvar IV, Fenno LE, Adhikari A, Lammel S, Mirzabekov JJ, Airan RD, Zalocusky KA, Tye KM, Anikeeva P, Malenka RC, Deisseroth K. Natural neural projection dynamics underlying social behavior. Cell. 2014; 157:1535–1551.

16. Harada K, Kamiya T, Tsuboi T. Gliotransmitter release from astrocytes: functional, developmental, and pathological implications in the brain. Front Neurosci. 2016; 9:499.

17. Kofuji P, Araque A. G-protein-coupled receptors in astrocyte-neuron communication. Neuroscience. 2021; 456:71–84.

18. Kofuji P, Araque A. Astrocytes and behavior. Annu Rev Neurosci. 2021; 44:49–67.

19. Kohls G, Perino MT, Taylor JM, Madva EN, Cayless SJ, Troiani V, Price E, Faja S, Herrington JD, Schultz RT. The nucleus accumbens is involved in both the pursuit of social reward and the avoidance of social punishment. Neuropsychologia. 2013; 51:2062–2069.

20. Krashes MJ, Koda S, Ye CP, Rogan SC, Adams AC, Cusher DS, Maratos-Flier E, Roth BL, Lowell BB. Rapid, reversible activation of AgRP neurons drives feeding behavior in mice. J Clin Invest. 2011; 121:1424–1428.

21. Le Merrer J, Detraux B, Gandía J, De Groote A, Fonteneau M, de Kerchove d’Exaerde A, Becker JAJ. Balance between projecting neuronal populations of the nucleus accumbens controls social behavior in mice. Biol Psychiatry. 2024; 95:123–135.

22. Manduca A, Servadio M, Damsteegt R, Campolongo P, Vanderschuren LJMJ, Trezza V. Dopaminergic neurotransmission in the nucleus accumbens modulates social play behavior in rats. Neuropsychopharmacology. 2016; 41:2215–2223.

23. Meadows SM, Palaguachi F, Jang MW, Licht-Murava A, Barnett D, Zimmer TS, Zhou C, McDonough SR, Orr AL, Orr AG. Hippocampal astrocytes induce sex-dimorphic effects on memory. Cell Rep. 43:114278.

24. Nedergaard M, Verkhratsky A. Artifact versus reality-how astrocytes contribute to synaptic events. Glia. 2012; 60:1013–1023.

25. Newman SW. The medial extended amygdala in male reproductive behavior. A node in the mammalian social behavior network. Ann N Y Acad Sci 1999; 877:242–257.

26. Noh K, Cho W-H, Lee B-H, Kim D-K, Kim Y-S, Park K, Hwang M, Barcelon E, Cho YK, Lee CJ, Yoon B-E, Choi S-Y, Park H-Y, Jun SB, Lee SJ. Cortical astrocytes modulate dominance behavior in male mice by regulating synaptic excitatory and inhibitory balance. Nat Neurosci. 2023; 26:1541–1554.

27. O’Connell LA, Hofmann HA. The vertebrate mesolimbic reward system and social behavior network: a comparative synthesis. J Comp Neurol. 2011; 519:3599–3639.

28. Ortinski PI, Reissner KJ, Turner J, Anderson TA, Scimemi A. Control of complex behavior by astrocytes and microglia. Neurosci Biobehav Rev. 2022; 137:104651.

29. Perea G, Navarrete M, Araque A. Tripartite synapses: astrocytes process and control synaptic information. Trends in Neurosci. 2009; 8:421–431.

30. Shan Q, Hu Y, Chen S, Tian Y. Nucleus accumbens dichotomically controls social dominance in male mice. Neuropsychopharmacology. 2022; 47:776–787.

31. Schmidt SNL, Fenske SC, Kirsch P, Mier D. Nucleus accumbens activation is linked to salience in social decision making. Eur Arch Psychiatry Clin Neurosci. 2019; 269:701–712.

32. Stanley DA, Adolphs R. Toward a neural basis for social behavior. Neuron. 2013; 80:816–826.

33. Walsh JJ, Christoffel DJ, Malenka RC. Neural circuits regulating prosocial behaviors. Neuropsychopharmacology. 2023; 48:79–89.

34. Wang Q, Kong Y, Wu D-Y, Liu J-H, Jie W, You Q-L, Huang L, Hu J, Chu H-D, Gao F, Hu N-Y, Luo Z-C, Li X-W, Li S-J, Wu Z-F, Li Y-L, Yang J-M, Gao T-M. Impaired calcium signaling in astrocytes modulates autism spectrum disorder-like behaviors in mice. Nat Commun. 2021; 12:3321.

35. Xu S, Jiang M, Liu X, Sun Y, Yang L, Yang Q, Bai Z. Neural circuits for social interactions: from microcircuits to input-output circuits. Front Neural Circuits. 2021; 15:768294.

36. Yu X, Taylor AMW, Nagai J, Golshani P, Evans CJ, Coppola G, Khakh BS. Reducing astrocyte calcium signaling in vivo alters striatal microcircuits and causes repetitive behavior. Neuron. 2018; 99:1170–1187.

37. Yu X, Moye SL, Khakh BS. Local and CNS-wide astrocyte intracellular calcium signaling attenuation in vivo with CalExflox mice. J Neurosci. 2021; 41:4556–4574.

